# A single dorsal vagal complex circuit mediates the aversive and anorectic responses to GLP1R agonists

**DOI:** 10.1101/2025.01.21.634167

**Authors:** Warren T. Yacawych, Yi Wang, Guoxiang Zhou, Shad Hassan, Stace Kernodle, Frederike Sass, Martin DeVaux, Iris Wu, Alan Rupp, Abigail J. Tomlinson, Zitian Lin, Anna Secher, Kirsten Raun, Tune Pers, Randy J. Seeley, Martin Myers, Weiwei Qiu

**Affiliations:** Departments of Internal Medicine, University of Michigan, Ann Arbor, MI USA; Department of Molecular and Integrative Physiology, University of Michigan, Ann Arbor, MI USA; Department of Metabolism and Endocrinology, National Clinical Research Center for Metabolic Diseases, The Second Xiangya Hospital, Central South University, Changsha, 410000, China; Zhejiang University-University of Edinburgh Institute, International Campus, Zhejiang University, Haining, China; Center for Basic Metabolic Research, University of Copenhagen, Copenhagen, Denmark; Department of Surgery, University of Michigan, Ann Arbor MI USA; Center for Adipocyte Signaling (ADIPOSIGN), University of Southern Denmark, Odense, Denmark; Department of Endocrinology, The Second Affiliated Hospital, School of Medicine, Zhejiang University, Hangzhou, China; Global Drug Discovery, Novo Nordisk A/S, Maløv, Denmark; Research and Early Development, Novo Nordisk A/S, Bagsværd, Denmark

**Keywords:** GLP1R, NTS, AP, Food Intake, Obesity

## Abstract

GLP-1 receptor agonists (GLP1RAs) effectively reduce feeding to treat obesity, although nausea and other aversive side effects of these drugs can limit their use. Brainstem circuits that promote satiation and that mediate the physiologic control of body weight can be distinguished from those that cause aversion. It remains unclear whether brainstem *Glp1r* neurons contribute to the normal regulation of energy balance and whether GLP1RAs control appetite via circuits distinct from those that mediate aversive responses, however. Hence, we defined roles for AP and NTS *Glp1r*-expressing neurons (AP^Glp1r^ and NTS^Glp1r^ neurons, respectively) in the physiologic control of body weight, the GLP1RA-dependent suppression of food intake, and the GLP1RA-mediated stimulation of aversive responses. While silencing non-aversive NTS^Glp1r^ neurons interfered with the physiologic restraint of feeding and body weight, restoring NTS^Glp1r^ neuron *Glp1r* expression on an otherwise *Glp1r*-null background failed to enable long-term body weight suppression by GLP1RAs. In contrast, selective *Glp1r* expression in AP^Glp1r^ neurons restored both aversive responses and long-term body weight suppression by GLP1RAs. Thus, while non-aversive NTS^Glp1r^ neurons control physiologic feeding, aversive AP^Glp1r^ neurons mediate both the anorectic and weight loss effects of GLP1RAs, dictating the functional inseparability of these pharmacologic GLP1RA responses at a circuit level.

## Introduction

Obesity and its complications, including type 2 diabetes, represent enormous challenges to personal well-being and public health. Lifestyle changes such as exercise and/or dieting generally fail to mediate consistent and prolonged effects on body weight^1^, however. While bariatric surgery can effectively treat obesity, it is irreversible and inaccessible to most patients^2^. The current generation of GLP-1 receptor (GLP1R) agonists (GLP-1RAs) produce sustained weight loss approaching that of bariatric surgery^3,4^, however-improving cardiometabolic health for many afflicted with obesity and diabetes.

Unfortunately, all GLP1RAs (including exenatide, liraglutide (Lira), semaglutide (Sema), dulaglutide, and the dual GIP/GLP1R agonist tirzepatide) promote nausea and other aversive gastrointestinal side effects in nearly two thirds of patients taking these medications^3,4^, and many patients discontinue therapy^5^. Thus, separating appetite suppression from the nauseating effects of GLP1-RAs could improve the treatment of obesity.

Recent evidence suggests that the brainstem neuron populations responsible for promoting the aversive, nausea-associated responses to gut malaise are distinct from those that mediate the satiating responses to feeding^6–8^. Indeed, while activating *Glp1r* neurons either in the area postrema (AP; AP^Glp1r^ cells) or the *nucleus tractus solitarius* (NTS; NTS^Glp1r^ cells) can dramatically reduce food intake, activating AP^Glp1r^ neurons (but not NTS^Glp1r^ neurons) promotes conditioned taste avoidance (CTA; a surrogate for nausea in rodents) ^6,9^. Similarly, while simultaneously ablating AP^Glp1r^ and NTS^Glp1r^ neurons abrogates the suppression of food intake and body weight by GLP-1RAs, inhibiting AP^Glp1r^ neurons alone suffices to block CTA formation in response to GLP1RAs^6,9^.

Hence, while only AP^Glp1r^ neurons can provoke (and are required to mediate) aversive responses, activating either AP^Glp1r^ and NTS^Glp1r^ neurons can suppress feeding. Thus, NTS^Glp1r^ neurons potentially represent an enticing pharmacologic target by which to provide durable weight loss while mitigating aversive gastrointestinal events.

While the use of chemogenetics has clearly defined the capabilities of AP^Glp1r^ and NTS^Glp1r^ neurons^6,9^; the pharmacologic sufficiency of each of these two circuits remains undefined for GLP1RA action. Additionally, the potential roles for AP^Glp1r^ and NTS^Glp1r^ neurons in the physiologic control of body weight remain unclear. Hence, we sought to define physiologic roles for NTS^Glp1r^ and AP^Glp1r^ neurons, as well as their contributions to the anorectic effects of GLP-1RAs.

## Results

### Distinct roles for NTS^Glp1r^ and AP^Glp1r^ neurons in the physiologic control of body weight and aversion

To determine the requirement for AP^Glp1r^ or NTS^Glp1r^ neurons in the physiologic control of energy balance, we injected AAV-DIO-TetTox-GFP (or control virus) into the AP or NTS of *Glp1r^Cre^* mice to mediate the Cre-dependent expression of tetanus toxin in AP^Glp1r^ or NTS^Glp1r^ (AP^Glp1r-TT^ and NTS^Glp1r-TT^ mice, respectively) silencing these neurons by preventing their release of neurotransmitters (Fig. 1A, B).

**Figure 1:**
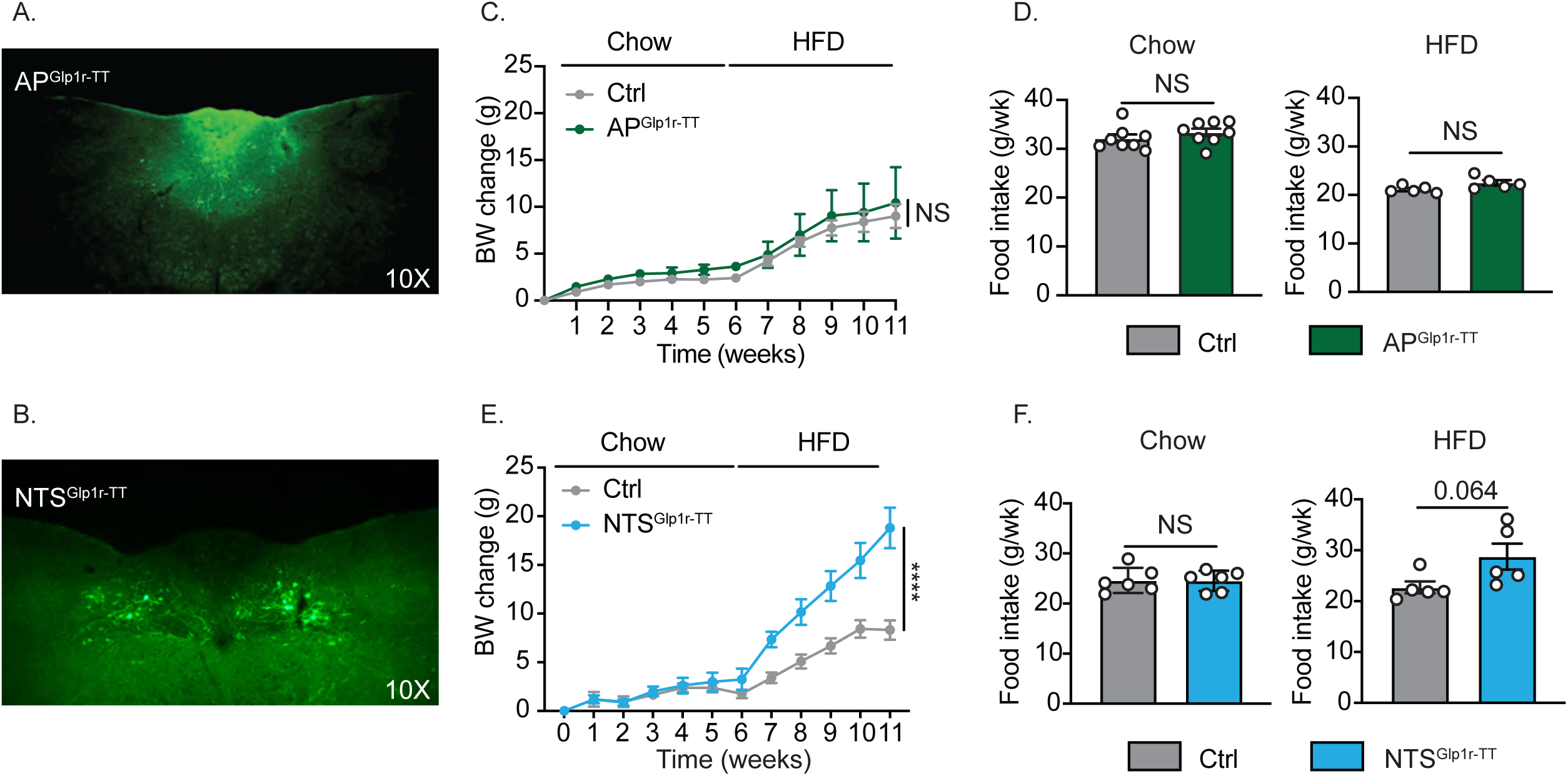
Silencing NTS^Glp1r^ neurons increases food intake and body weight during HFD feeding. Examples of GFP-IR (green) in the DVC of AP^Glp1r-TT^ (A) and NTS^Glp1r-TT^ (B) mice (images acquired at 10X magnification). (C) Body weight change (grams) following surgery for AP^Glp1r-TT^ (n=4) and control (Ctrl;n=10) mice on chow for 6 weeks followed by HFD for 5 weeks. (D) Weekly food intake for AP^Glp1r-TT^ (n=5-8) and control (n=5-8) mice during the final week of chow (left) or HFD (right) feeding. (E) Body mass gained following surgery for NTS^Glp1r-TT^ (n=5) or Ctrl (n=6) mice on chow for 6 weeks followed by HFD for 5 weeks. (F) Food intake of NTS^Glp1r-TT^ (n=5-6) or control (n=6) mice during the final week of chow (left) or HFD (right) feeding. All data presented as mean ± SEM. Repeated measures two-way ANOVA Sidaks multiple comparisons for C, E. Unpaired two-tailed t-test for D, F. NS p>0.05, *p<0.05, **p<0.01, ***p<0.001, ****p<0.0001.

Following surgery, we measured food intake and body weight in AP^Glp1r-TT^ (Figure 1 C-D) and NTS^Glp1r-TT^ (Fig. 1E-F) mice during 6 weeks on normal chow diet followed by another five weeks with high fat diet (HFD) feeding. Silencing AP^Glp1r^ neurons failed to alter food intake or body weight compared to controls on either diet (Fig. 1C-D). In contrast, while silencing NTS^Glp1r^ neurons did not detectably alter food intake or body weight compared to controls in chow-fed NTS^Glp1r-TT^ mice, these animals tended to eat approximately 25% more food and gained almost twice as much weight as controls over the six weeks of HFD-feeding (Fig. 1E-F).

Hence, the endogenous activity of NTS^Glp1r^ neurons (but not AP^Glp1r^ neurons) mediates the normal restraint of food intake in HFD-fed mice and contributes to the physiologic control of body weight. Furthermore, we confirmed that chemogenetically activating AP^Glp1r^ neurons, but not NTS^Glp1r^ neurons, promote aversion (Supplemental Fig. 1A-D) ^6,9^, and that activating either population sufficed to mediate the multiday suppression of food intake and body weight (Supplemental Fig. 1E-H). These findings suggest that the GLP1RA-dependent activation of AP^Glp1r^ neurons should suppress feeding and promote CTA formation, while GLP1RA action on NTS^Glp1r^ neurons should non-aversively suppress food intake.

### Roles for GLP1R on AP^Glp1r^ cells in GLP1RA-mediated aversive and feeding responses

To assess roles for AP GLP1R in the anorectic effects of Lira, we stereotaxically injected AAV-Cre into the AP of *Glp1r^fl/fl^* mice (generating AP^Glp1r-KO^ mice predicted to lack *Glp1r* in the AP); an injection of AAV-tdTomato generated control AP^Ctrl^ mice (Supplemental Fig 2A-B). We administered Lira to these mice at the onset of the dark cycle and assessed food intake over the subsequent 24 hours (Supplemental Fig 2C). While the Lira-stimulated suppression of food intake was attenuated in AP^Glp1rKO^ animals compared to controls during the first 4 hours of treatment, this effect was no longer detectable at 24 hours. The latter finding is consistent with the prior observation suggesting that neither the ablation of *Glp1r* in the AP nor the chemogenetic inhibition of AP^Glp1r^ neurons alters GLP1RA-dependent food intake over 24 hours^6^.

While we found that Lira promoted CTA formation in AP^Glp1r-KO^ mice, this response was attenuated and more variable compared to AP^Ctrl^ mice (Supplemental Fig 2D), consistent with the previously demonstrated role for AP^Glp1r^ neurons in mediating aversive responses to GLP1RA treatment^9^. The variability in the magnitude of CTA formation to Lira exhibited by AP^Glp1r-KO^ mice led us to surmise that AP^Glp1r-KO^ mice might exhibit incomplete ablation of AP *Glp1r*, however. Indeed, we found substantial Lira-induced FOS-immunoreactivity (-IR) in the AP of AP^Glp1r-KO^ mice (Supplemental Fig 2E-F), suggesting the likely persistence of AP *Glp1r* expression in these mice. This presumably reflects the difficulty of ablating enough AP *Glp1r* expression with AAV-CRE injection to disrupt AP GLP1RA action while restricting viral transduction enough to avoid spillover into the NTS.

Thus, although our results demonstrate that AP GLP1R plays a previously unreported role in the acute (up to 4 hours) GLP1RA-mediated suppression of food intake, the incomplete ablation of GLP1R in the AP of AP^Glp1r-KO^ mice renders it impossible to fully understand the roles for AP GLP1R in GLP1RA responses using this mouse model.

### Site-specific reactivation of *Glp1r* expression in the DVC

To isolate the contributions of AP and NTS GLP1R to GLP1RA action, we thus turned to the *Glp1r^LoxTB^*allele, in which a LoxP-flanked transcription blocking cassette lies upstream of the fourth exon of the endogenous *Glp1r* gene^10^. Cre removes the transcription blocker from the *Glp1r^LoxTB^* allele, reactivating native *Glp1r* expression on an otherwise *Glp1r*-null background.

While we have substantial experience with comprehensively transducing the NTS without viral leakage into the nearby AP, we theorized that transducing the AP with virus to sufficiently reactivate AP *Glp1r* expression without promoting NTS *Glp1r* expression could present challenges similar to those we observed in the AP^Glp1r-KO^ mice (Supplemental Fig. 2). Hence, we set out to identify a marker gene specific to AP^Glp1r^ neurons that we could use as a Cre driver to permit the genetic reactivation of *Glp1r* expression from the *Glp1r^LoxTB^* allele in the AP specifically.

We crossed *Glp1r^Cre^* onto the *Rosa26^tm5.1(CAG-Sun1-sfGFP)Nat^* background to promote the Cre-mediated expression of the nuclear envelope-labeling GFP-SUN1 fusion protein in *Glp1r* neurons^11,12^. We dissected the hypothalamus, DVC, and nodose ganglion from these mice, used FACS sorting to enrich for GFP-containing nuclei, and subjected the resultant nuclei to single nucleus RNA sequencing (snRNA-seq) (Supplemental Fig 3A).

This analysis identified 78 clusters of *Glp1r* neurons (Supplemental Fig 3B). To identify AP^Glp1r^ and NTS^Glp1r^ neurons, we integrated our dataset with previous snRNA-seq atlases of the hypothalamus^13^, DVC^14^, and nodose^15^. While examining cell type-specific gene expression for these populations identified no useful marker gene for NTS^Glp1r^ neurons, this analysis revealed strong *Pirt* expression in AP^Glp1r^ neurons (Supplemental Fig 3C). While the previously identified cluster of GABAergic hypothalamic arcuate nucleus (ARC) neurons marked by *Tbx19* expression^10,16^ also contained a few *Pirt-*expressing cells, the previous finding that *Glp1r* in GABA neurons or in the hypothalamus minimally impacts GLP1RA action^17,18^ suggests these cells are unlikely to contribute GLP1RA-mediated food intake control.

We thus crossed the previously-described *Pirt^Cre^* allele^19^ onto the *Glp1r^LoxTB^* background to generate *Pirt^Cre^;Glp1r^LoxTB/LoxTB^* mice predicted to express *Glp1r* specifically in the AP (AP^Glp1r-Re^ mice). We perfused some of these mice to examine the distribution of GLP1R-IR in these animals. As expected, we detected no GLP1R in *Glp1r^LoxTB/LoxTB^* (Glp1r^KO^) mice (Fig. 2A). In contrast, AP^Glp1r-Re^ mice displayed similar AP GLP1R-IR as wild-type controls (Figure 2B, D). We detected no GLP1R-IR in the ARC (Supplemental Fig 3E) or elsewhere in the brain (data not shown) of AP^Glp1r-Re^ mice, while GLP1R-IR detection in animals with the wild-type *Glp1r* allele (Glp1r^WT^) matched its expected distribution in the ARC (Supplemental Fig. 3D) and throughout the brain (not shown). Furthermore, AP^Glp1r-Re^ animals demonstrated restored Lira-dependent CTA formation (Supplemental Fig 3H) and FOS accumulation in CGRP neurons of the parabrachial nucleus (PBN) (Supplemental Fig 3I-J), whose activity is sufficient and required for CTA formation^20^. Thus, *Pirt^Cre^* specifically and robustly restores AP GLP1R signaling.

**Figure 2:**
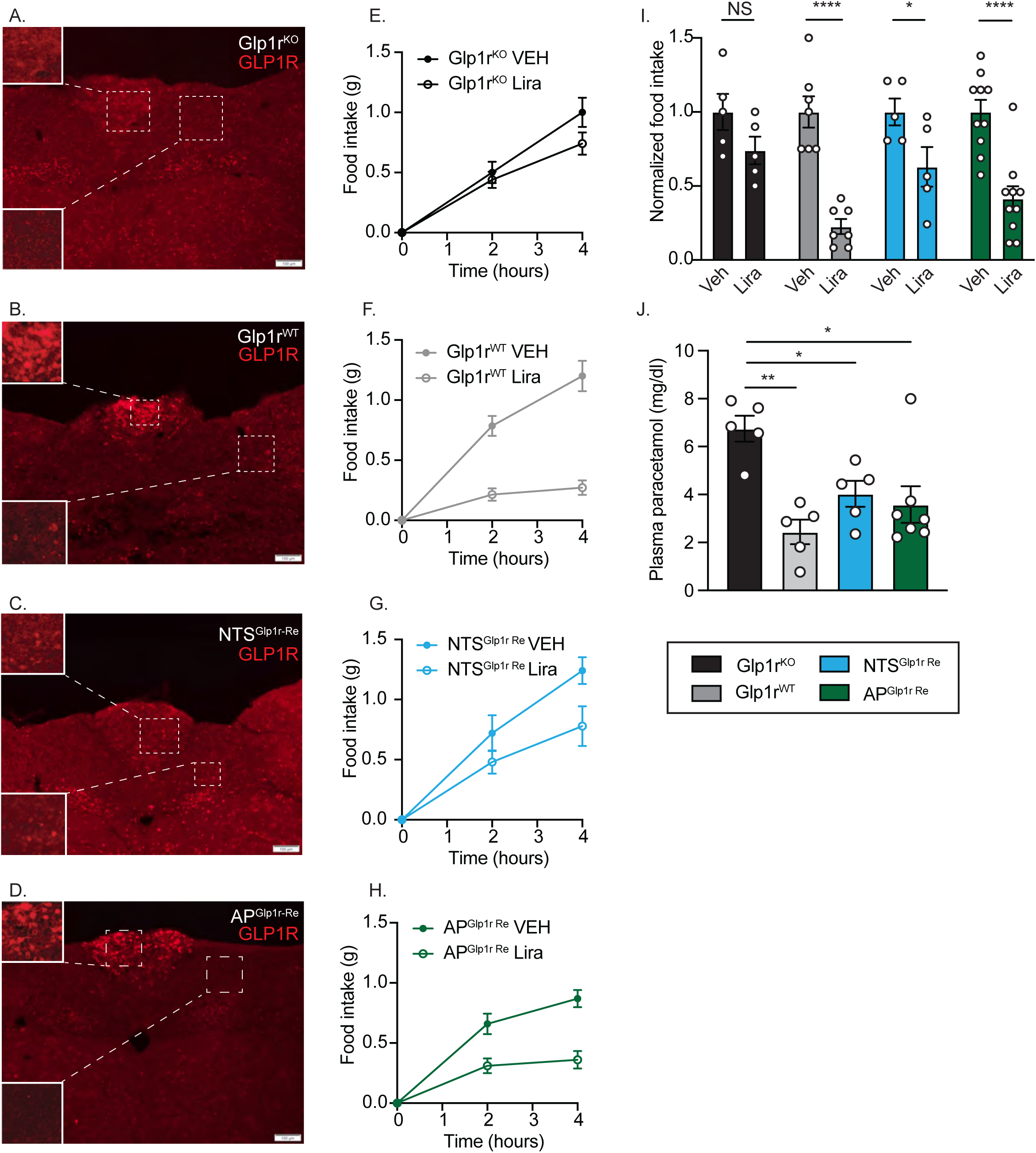
Acute reduction of food intake and gastric emptying by Lira in AP^Glp1r-Re^ and NTS^Glp1r-Re^ animals. A-D: Representative images of GLP1R-IR (red) in the DVC of control Glp1r^KO^ (A; n=5), Glp1r^WT^, (B; n=7), NTS^Glp1r-Re^ (C; n=5), and AP^Glp1r-Re^ (D; n=10) animals. Scale bar=100 um). E-H: HFD intake during the first four hours of the dark cycle for the indicated lines following saline (Veh) or Lira (400 ug/kg, IP) injection. (I) Food intake normalized to average 4-hour food intake in Veh-treated animals over the first four hours of the dark cycle from (E-H). (J) Plasma paracetamol concentrations 15 minutes after gavage of paracetamol in saline for Lira-treated Glp1r^KO^ (n=5), Glp1r^WT^ (n=5), NTS^Glp1r-Re^ (n=5), and AP^Glp1r-Re^ (n=7) animals. All graphs: mean ± SEM are shown. Repeated measures two-way ANOVA with Sidak’s multiple comparisons for E-H. Two-way ANOVA with Tukey’s multiple comparisons for I. One-way ANOVA with Tukey’s multiple comparisons for J. NS p>0.05, *p<0.05, **p<0.01, ***p<0.001, ****p<0.0001.

As expected, *Glp1r^LoxTB/LoxTB^* mice that had been injected in the NTS with AAV-Cre-GFP (NTS^Glp1r-Re^ mice; Supplemental Fig. 3F) displayed GLP1R-IR in the NTS (Fig 2C). While GFP staining revealed no detectable leak of AAV-Cre-GFP into the AP of NTS^Glp1r-Re^ mice (Supplemental Fig 3F-G), we observed small numbers of GLP1R-IR soma in the AP of these animals (Fig 2C). Because most serotypes of AAVs undergo varying degrees of uptake from axonal terminals^21–23^ we speculate that these few GLP1R-IR AP neurons in the NTS^Glp1r-Re^ mice result from the uptake of a few AAV-Cre-GFP genomes into small numbers of NTS-projecting AP^Glp1r^ neurons.

To define roles for *Glp1r* in the AP and NTS for the effects of acute Lira treatment on food intake and gastric emptying, we administered Lira or vehicle to Glp1r^WT^, Glp1r^KO^, AP^Glp1r-Re^, and NTS^Glp1r-Re^ mice at the onset of the dark cycle and measured food intake over the subsequent 4 hours (Fig 2E-H). While Lira failed to significantly decrease food intake in Glp1r^KO^ mice, it suppressed food intake in Glp1r^WT^ animals by approximately 80% during the first 4 hours of the dark cycle. Lira also suppressed food intake in NTS^Glp1r-Re^ and AP^Glp^^1-Re^ animals, although the suppression of food intake in NTS^Glp1r-Re^ mice (approximately 35%) was less pronounced than in AP^Glp1r^ ^Re^ mice (approximately 60%) (Fig 2I).

Furthermore, Lira decreased the uptake of gavaged paracetamol into the bloodstream (a measure of gastric emptying rate) in Glp1r^WT^, NTS^Glp1r-Re^ and AP^Glp1r-Re^ mice (Fig 2J). These data suggest that GLP1RAs can act via GLP1R in the AP and in the NTS to rapidly slow gastric emptying and suppress feeding at the onset of the dark cycle.

### AP GLP1R mediates the GLP1RA-dependent long-term suppression of food intake and body weight

Some *Glp1r*-expressing neurons, like those in the nodose ganglion and the dorsomedial hypothalamic nucleus (DMH), mediate acute but not long-term food intake suppression^10,18,24^. We thus determined the ability of AP^Glp1r^ and NTS^Glp1r^ neurons to mediate long-term effects on energy balance by examining the long-term feeding and body weight responses of NTS^Glp1r-Re^ and AP^Glp1r-Re^ mice to GLP1RA treatment (Figures 3-4).

**Figure 3:**
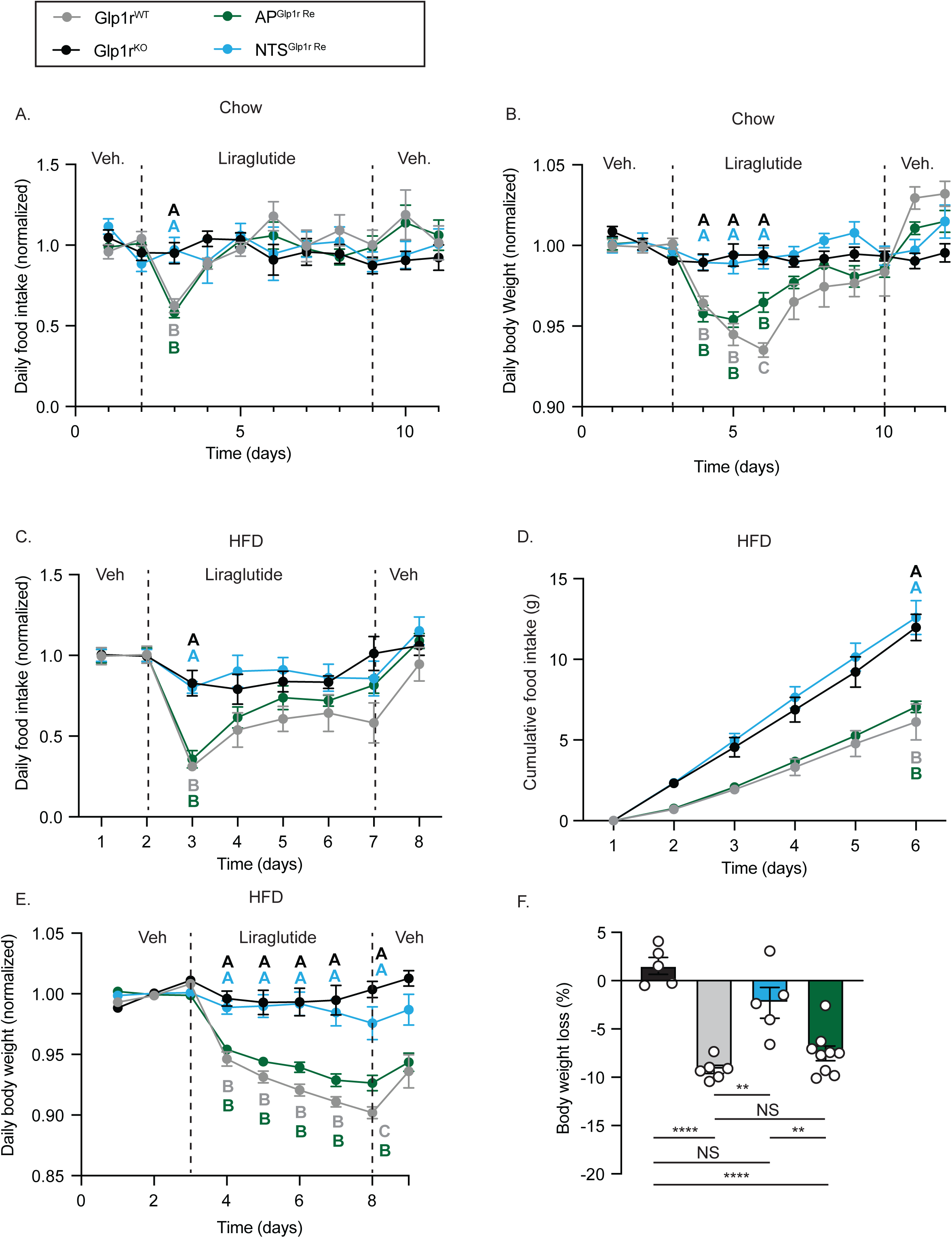
Lira reduces food intake and body weight in AP^Glp1r-Re^ but not NTS^Glp1r-Re^ mice. The indicated strains of mice were injected with vehicle (Veh) for 2 days, Lira for 5-7 days, and Veh again for two days; food intake and body weight were measured daily. (A-B) Food intake (A) and body weight (B) normalized to baseline for chow-fed Glp1r^KO^ (n=6), Glp1r^WT^ (n=7), NTS^Glp1r-Re^ (n=5), and AP^Glp1r-Re^ (n=10) mice. (C-F) Daily food intake over the course of the experiment. Daily food intake normalized to baseline for Veh or Lira treated mice and (D) cumulative food intake over the course of Lira treatment. Daily body weight normalized to baseline (E) and total percent change in body weight (F) during Lira treatment for HFD-fed Glp1r^KO^ (n=5), Glp1r^WT^ (n=6), NTS^Glp1r-Re^ (n=5), and AP^Glp1r-Re^ (n=9) mice. All graphs: mean +/- SEM shown. Repeated measures two-way ANOVA with Tukey’s multiple comparisons for A-E. One-way ANOVA with Tukey’s multiple comparisons for F. A-F: Groups (indicated by letter matching color of line) with different letters are different at the indicated time point, p<0.05. F: NS p>0.05, * =p<0.05, **= p<0.01, *** =p<0.001, **** = p<0.0001.

**Figure 4:**
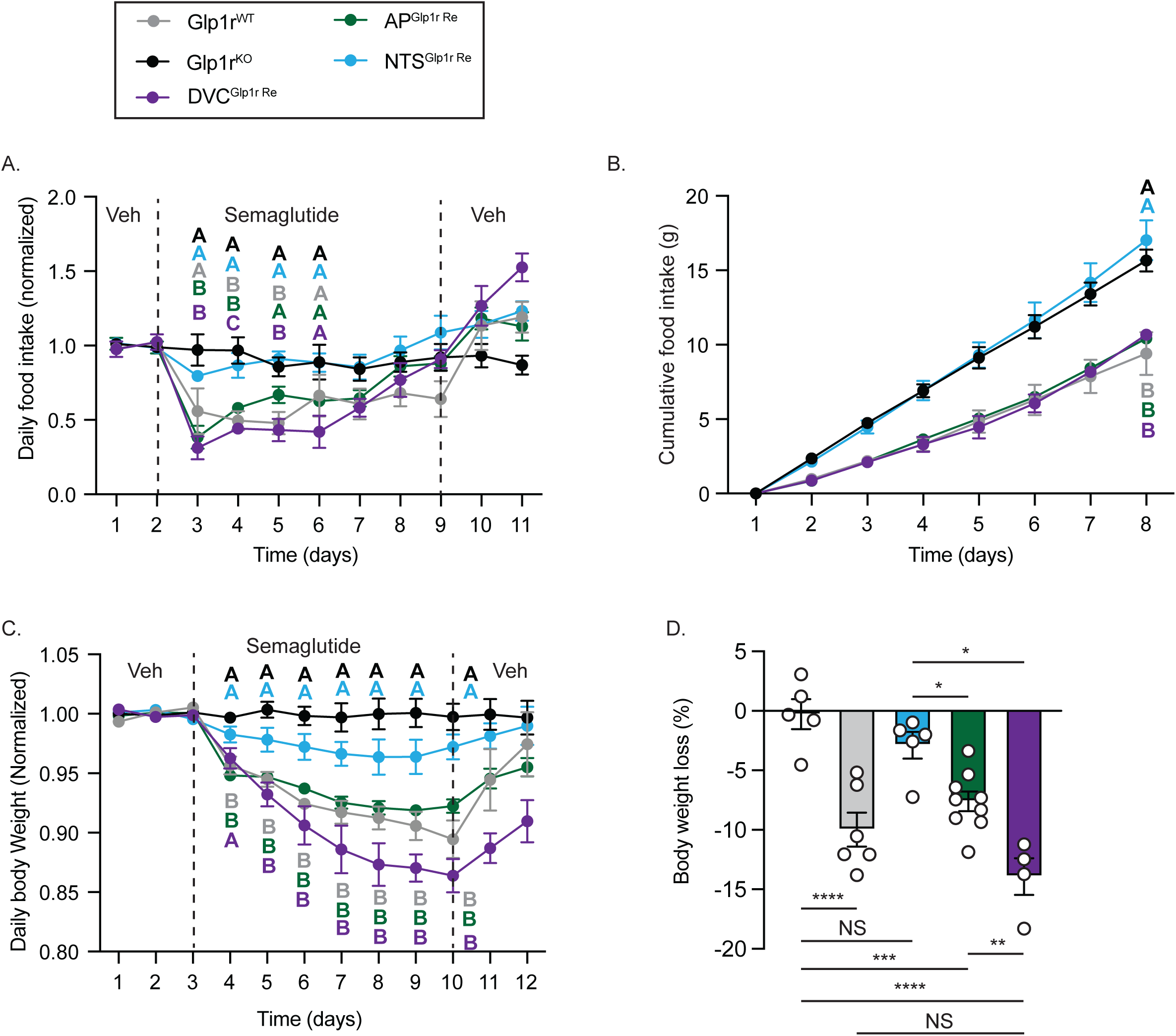
Effects of Sema on food intake and body weight in AP^Glp1r-Re^, NTS^Glp1r-Re^, and DVC^Glp1r-Re^ animals. Animals were treated with vehicle (Veh) for two days, Sema for 7 days, and Veh for 2 additional days. Food intake and body weight were measured once daily. Shown are daily food intake over the course of the experiment (A), cumulative food intake during Sema treatment (B), daily body weight (C) and overall change in body weight during Sema treatment for HFD-fed Glp1r^KO^ (n=5), Glp1r^WT^ ( n=6), NTS^Glp1r-Re^ (n=5), AP^Glp1r-Re^ (n=9), and DVC^Glp1r-Re^ (n=4) animals. A and C are normalized to baseline food intake measures. All graphs: mean +/- SEM shown. Repeated measures two-way ANOVA with Tukey’s multiple comparisons for A-C. One-way ANOVA with Tukey’s multiple comparisons for D. A-C: Groups (indicated by letter matching color of line) with different letters are different at the indicated time point, p<0.05. D: NS=p<0.05. F: NS p>0.05, * =p<0.05, **= p<0.01, *** =p<0.001, **** = p<0.0001.

We initially treated chow-fed Glp1r^WT^, Glp1r^KO^, NTS^Glp1r-Re^, and AP^Glp1r-Re^ animals with vehicle for 2 days, Lira for seven days, and then with vehicle for an additional two days. We monitored food intake and body weight each day at the time of treatment, one hour prior to the onset of the dark cycle. Interestingly, while Lira decreased food intake by approximately 40% in lean Glp1r^WT^ animals over the first day of treatment, food intake in these animals subsequently returned towards baseline levels despite ongoing treatment (Figure 3A). Similarly, Lira decreased body weight in Glp1r^WT^ mice for the first 3 days of treatment only, after which body weight began to increase toward baseline (Figure 3B).

While Lira provoked no changes in food intake and body weight in Glp1r^KO^ mice, the suppression of food intake in AP^Glp1r-Re^ mice mirrored that of Glp1r^WT^ animals, although body weight began to return to normal more rapidly in the AP^Glp1r-Re^ mice than in Glp1r^WT^ mice. Surprisingly, we detected no alterations in food intake or body weight in NTS^Glp1r-Re^ animals. While these findings suggest AP GLP1R suffices to mediate the majority of long-term GLP1RA action on feeding and body weight, they also indicate that lean, chow-fed animals represent a suboptimal model for the study of GLP1RA action.

Given the strong suppression of feeding and body weight in response to the chemogenetic activation of NTS^Glp1r^ neurons and the important role that NTS^Glp1r^ neurons play in the physiologic control of energy balance, the lack of NTS^Glp1r-Re^ animals to respond to GLP1RAs was somewhat surprising. Because of the limited magnitude and duration of the Lira response in lean, chow-fed animals, and the HFD-specific role for NTS^Glp1r^ neurons in the physiologic control of energy balance, we reasoned that NTS^Glp1r^ neurons might play more prominent roles in obese HFD-fed animals. We fed HFD to all the groups of mice for several weeks, and then re-assessed their response to Lira during HFD feeding (Figure 3C-F).

While Lira did not alter food intake or body weight in obese Glp1r^KO^ mice, Lira decreased food intake and body weight across the entire 5-day treatment period in the obese Glp1r^WT^animals and promoted similar effects in AP^Glp1r-Re^ mice (Figure 3C-F). Strikingly, however, Lira once again failed to alter food intake and body weight in obese NTS^Glp1r-Re^ animals. Thus, restoring AP *Glp1r* expression alone produced a Lira response similar to that of GLP1R^WT^ mice, while reactivating *Glp1r* expression in the NTS failed to restore any long-term Lira responses. These data suggest that NTS GLP1R contributes little to the catabolic response to Lira, while GLP1R in AP^Glp1r^ neurons suffices to mediate most long-term Lira action in the suppression of feeding and body weight.

While Lira durably suppresses appetite and body weight in humans and rodents, Sema promotes larger reductions in food intake and body weight^25^. Mechanisms for this increased efficacy may include its longer half-life/duration of action and/or improved penetration beyond the blood-brain barrier (BBB) into deep brain regions than Lira^26^. Because the AP resides outside the BBB and is freely accessible to circulating peptides, while the NTS lies behind the BBB^27^, we thus examined the response of HFD-fed AP^Glp1r-Re^ and NTS^Glp1r-Re^ mice to Sema (Fig 4). Furthermore, to determine whether the restoration of NTS *Glp1r* expression might augment the response to GLP1RA treatment in AP^Glp1r-Re^ mice, we injected AAV-Cre-GFP into the NTS of AP^Glp1r-Re^ animals (generating DVC^Glp1r-Re^ mice) and examined their response to Sema, as well.

As with Lira, Sema decreased food intake (Fig 4A-B) and body weight (Fig 4C-D) in AP^Glp1r-Re^ and Glp1r^WT^ HFD-fed animals. Also as we observed with Lira, the body weight and food intake responses of NTS^Glp1r-Re^ animals to Sema were not significantly different than for Glp1r^KO^ mice. Interestingly, however, Sema promoted a slightly greater decrease in food intake and body weight in DVC^Glp1r-Re^ mice than in AP^Glp1r-Re^ animals. Thus, GLP1RA action via NTS^Glp1r^ neurons in isolation fails to significantly suppress feeding or body weight, but these neurons may mediate a small synergistic effect when pharmacologically targeted in tandem with AP^Glp1r^ neurons. AP^Glp1r^ neurons mediate most GLP1RA-induced weight loss, however.

## Discussion

Here, we reveal distinct physiologic and pharmacologic roles for brainstem *Glp1r-*expressing circuits in the control of feeding, body weight, and aversion. While non-aversive NTS^Glp1r^ neurons are required for the physiologic long-term restraint of food intake and body weight, these cells fail to mediate weight loss when targeted by GLP1RAs. In contrast, AP^Glp1r^ neurons not only mediate the aversive nausea-like responses to GLP1RAs but also mediate the majority of the anorectic and weight loss effects of these drugs.

These results suggest that the *Glp1r*-expressing circuits important for nausea (AP^Glp1r^ neurons) and the normal physiologic control of satiety (NTS^Glp1r^ neurons) are distinct, consistent with the finding that gastrointestinal irritants activate only AP^Glp1r^ neurons while NTS^Glp1r^ neurons can respond to nutrient infusion to the gut^6^. Our results also suggest that a single site of action in the brainstem (AP^Glp1r^ neurons) mediates both the aversive and weight loss effects of GLP1RA pharmacotherapy, however. Hence, it would be difficult to separate the aversive response from the anorectic effects of GLP1RAs at the level of the *Glp1r*-expressing cell types/circuits on which they act.

Given the strong anorectic response promoted by the chemogenetic activation of NTS^Glp1r^ neurons and the importance of these neurons for the physiologic control of energy balance, we were initially surprised by the failure of direct GLP1RA action on NTS^Glp1r^ neurons to suppress food intake and energy balance in a sustained manner. Blocking the overall action of a particular set of neurons (e.g., by inhibiting or silencing them) can interfere with a host of functions above and beyond those mediated by a single receptor on that neuron, however. Indeed, our findings reveal that NTS^Glp1r^ neurons restrain food intake and body weight in response to undefined endogenous signals, but not in response to GLP1RAs.

Explanations for this failure of NTS^Glp1r^ neurons to mediate the GLP1RA-stimulated suppression of food intake and body weight could include insufficient blood-brain barrier (BBB) penetration for the GLP1RAs tested (the NTS resides behind the BBB while the AP lies outside it)^27^. The finding that fluorescent Sema accumulates in the NTS and other structures behind the BBB^26^ argues against a problem with GLP1R access to NTS^Glp1r^ neurons, however. Furthermore, Ca^+2^ imaging reveals the GLP1RA-mediated activation of NTS^Glp1r^ neurons *in vivo*^6^ and GLP1RA treatment of NTS^Glp1r-Re^ animals delays gastric emptying and promotes short-term effects on food intake, suggesting that GLP1RAs are able to rapidly access and modulate the activity of NTS^Glp1r^ cells.

It is possible the NTS lacks sufficient GLP1R expression to sustain periods of increased activity, however. Indeed, we observe that GLP1R-IR is less robust in the NTS than the AP and the magnitude of the short-term NTS^Glp1r^-mediated responses to GLP1RA tend to be smaller than those mediated by AP^Glp1r^ neurons. NTS^Glp1r^ cells are also fewer in number than AP^Glp1r^ neurons. Indeed, GLP1RAs poorly promote FOS accumulation in NTS^Glp1r^ neurons^17^. Hence, the relatively low GLP1R content on the relatively few NTS^Glp1r^ neurons may not promote sufficient signaling by these cells to strongly and durably suppress feeding. The notion that GLP1RAs may access NTS^Glp1r^ neurons but that this direct GLP1RA→NTS^Glp1r^ neuron action fails to mediate substantial long-term changes in feeding and energy balance predicts that the generation of GLP1RAs with improved BBB penetrance (or other pharmacodynamic alterations) might do little to augment the strength of the NTS^Glp1r^ neuron-mediated response to such compounds. Rather, NTS^Glp1r^ neurons (or other NTS neurons) may represent better candidates for obesity treatment via agonists for other receptors that they express at higher levels.

Our inability to completely ablate *Glp1r* from all AP^Glp1r^ neurons by stereotaxic injection suggests that we cannot rule out the possibility that the failure to identify roles in the physiologic restraint of feeding by AP^Glp1r^ neurons might result from incomplete transduction and silencing of these cells in AP^Glp1r-TT^ mice. The notion that AP^Glp1r^ neurons do not contribute to the normal control of body weight is consistent with the finding that lesioning the AP does not increase long term body mass on chow or HFD, however^28^.

While the intra-NTS injection of AAV-Cre-GFP promoted GLP1R-IR in a few AP^Glp1r^ neurons (in addition to restoring GLP1R-IR in the NTS), the finding that NTS *Glp1r* expression augments the AP^Glp1r^ neuron-mediated suppression of food intake and body weight by GLP1RAs suggests the potential existence of NTS^Glp1r^-dependent GLP1RA effects that are additive to those mediated by AP GLP1R. The persistence of this additive effect of GLP1RA action via NTS^Glp1r^ neurons during subchronic treatment also argues against tachyphylaxis as a mechanism that might limit the NTS^Glp1r^ neuron-mediated GLP1RA response.

We previously showed that silencing NTS *Calcr* cells, which map to the same bioinformatically-defined neuron population as NTS^Glp1r^ neurons, interferes with food intake suppression by exendin-4^8^. Hence, NTS^Glp1r^ neurons might function similarly to NTS *Calcr* cells, acting in a downstream neural system that participates in the appetite-suppressing effects of AP^Glp1r^ neurons, such that modestly increasing the activity/excitability of NTS^Glp1r^ neurons could augment the anorectic signal from AP^Glp1r^ cells. We cannot exclude the possibility that the reactivation of *Glp1r* expression in small numbers of AP^Glp1r^ neurons mediate the modest GLP1RA-dependent effects observed in Glp1r^NTS-Re^ mice, however. In either case, our data demonstrate that GLP1R in the NTS represents a minor pathway for overall food intake control by GLP1RAs.

Roles in GLP1RA-dependent feeding suppression have been suggested for *Glp1r*-expressing neurons in the lateral septum (LS; LS^Glp1r^ neurons)^29^ or DMH (DMH^Glp1r^ neurons)^24,30^. While it is possible these circuits influence aspects of appetite other than suppressing overall food intake (i.e., reward, palatability, post-ingestive feedback), our data demonstrate the importance of the AP for the GLP1RA-mediated control of food intake over the long term, however. Indeed, the Sema-stimulated long-term transcriptional response is much stronger in the AP than in the NTS, LS, or DMH^14^.

Furthermore, while GABAergic DMH^Glp1r^ neurons directly innervate and inhibit orexigenic ARC *Agrp* neurons^30^, restoring DMH *Glp1r* permits only the short-term suppression of food intake by GLP1RAs, not the long-term suppression of food intake and body weight^10,24^.

Additionally, ablating *Glp1r* in the LS only slightly decreases the anorectic response to GLP1RAs^29^. While (like NTS^Glp1r^ neurons) LS^Glp1r^ neurons may restrain physiologic feeding, the extent to which direct responses to GLP1RA treatment by LS^Glp1r^ neurons mediate these effects remains undefined. While our study identifies the AP as a major site of action for GLP1RA-induced weight loss, it remains possible other *Glp1r-*expressing circuits coordinate responses to GLP1RAs that are poorly contextualized by measuring overall food consumption. Hence, it will be interesting to examine the sufficiency of GLP1R in LS^Glp1r^ neurons to mediate GLP1RA effects, along with the ability of LS^Glp1r^ and DMH^Glp1r^ cells to augment AP^Glp1r^-mediated responses to GLP1RAs.

Overall, our results demonstrate that AP^Glp1r^ neurons suffice to mediate the long-term anorectic responses to GLP1RAs. Because non-aversive NTS^Glp1r^ neurons contribute little to long-term GLP1RA action, the finding that AP GLP1R mediates aversion as well as most of the long-term reduction of food intake and body weight by GLP1RAs suggests the likely inability to separate the anorectic and aversive responses of GLP1RAs by selectively targeting GLP1R in different brainstem areas or cell types. Several snRNA-seq analyses demonstrate the existence of at multiple distinct *Glp1r* expressing neuron populations within the AP^9,14,31^, however. Hence, it remains possible that the suppression of food intake and body weight might be mediated by AP *Glp1r* neuron subtypes different than those that mediate aversion. Consequently, it will be important to define roles for each population of AP^Glp1r^ neurons in GLP1RA action. If distinct AP^Glp1r^ neuron subtypes mediate the aversive versus appetite-suppressing effects of GLP1RAs, this could provide a potential neural mechanism by which to separate the salutary and aversive effects of GLP1RAs.

## Methods

### Animals

Mice for AP stereotaxic injections were bred and housed in animal colonies within the animal facility at Zhejiang University. All animal experiments were performed according to the protocols approved by the Institutional Animal Care and Use Committee (IACUC) of Zhejiang University (approval number: 23311) in accordance with the institutional guidelines.

All other animals were bred and housed in the Unit for Laboratory Animal Medicine at the University of Michigan. These mice and the procedures performed were approved by the University of Michigan IACUC and in accordance with Association for the Assessment and Approval of Laboratory Animal Care and National Institutes of Health guidelines.

Mice were housed in specific pathogen free (SPF), temperature controlled (22° C) rooms with 12-hour light and dark cycles. Mice were provided with food (Purina Lab Diet 5LOD, unless otherwise specified) and water *ad libitum* (except as noted below) with daily health checks performed by qualified animal personnel.

We purchased male and female C57BL/6 mice for experiments and breeding at the University of Michigan from Jackson Laboratories. *Glp1r^Cre^* (Jax cat #029283), *Glp1r^Stop-Flo^*^x 10^, *Glp1r^Flox^* ^18^, and *Pirt^Cr^*^e^ ^19^ animals have been previously described. *Glp1r^Cre^*, *Glp1r^Flox^*, and *Glp1r^Stop-Flox^* were propagated by intercrossing homozygous mice of the same genotype. We crossed *Pirt^Cre/+^; Glp1r^Stop-Flox/Stop-Flox^*animals with homozygous *Glp1r^Stop-Flox/Stop-Flox^* mice to generate *Pirt^Cre/+^_;_ Glp1r^Stop-Flox/Stop-Flox^* (AP^Glp1r-Re^) and *Glp1r^Stop-Flox/Stop-Flox^* (Glp1r^KO^) animals for study.

For the AP stereotaxic injections *Glp1r^Cre^* mice were purchased from Shanghai Model Organisms Center, Inc. (Shanghai, China) (Cat No: NM-KI-200134); *Glp1r^Flox^*mice were purchased from GemPharmatech (Nanjing, China) (Cat No: T005818).

For all studies, animals were processed in order of ear tag number, which was randomly assigned at the time of tailing (before genotyping). All mouse models (with the exception of purchased C57BL/6 mice) were on the segregating C57BL/6;SJL;129 genetic background; sibling controls were used throughout. Male and female mice were used for all experiments where a single sex was not indicated; no sex differences in phenotype were detected.

### Viral Reagents and Stereotaxic Injections

AAV-Cre-GFP, AAV-GFP, AAV-DIO-hM3Dq-mCherry, and AAV-DIO-TetTox-GFP were as previously described^8^ and were prepared as AAV8 serotype viruses by the University of Michigan Viral Vector Core.

AAV-hSyn-tdTomato (S0266-8), AAV-hSyn-Cre-mCherry (AAV-Cre-mCherry, S0702-8), AAV-hSyn-DIO-hM3Dq-mCherry (S0192-8) and AAV-CAG-DIO-Tettox-EGFP (S0235-8) for AP injections were purchased from Shanghai Taitool Bioscience (Shanghai, China) specifically for AP injections.

For injection, following the induction of isoflurane anesthesia and placement in a stereotaxic frame, the skulls of adult mice were exposed. Neck muscles were carefully cut until the brainstem was exposed.

For NTS injection, the fourth ventricle was set as reference point for injection. The distance across the ventricle M/L=0.25mm, and D/V=0.5 was used. After the reference was determined, a guide cannula with a pipette injector was lowered into the NTS and 50 nl of virus was injected using a World Precision Instrument microinjector at a rate of 5-30 nl/minute.

For AP injection, the obex was set as a reference location, the dura above brain tissue was opened to locate the injection site with a coordinate: D/V: -0.2 mm; A/P: +0.25 mm; M/L: 0 mm. 30 nl AAVs under 1 nl/sec rate using a brain stereotaxic injection apparatus (R480 glass micro-glass injection pump, RWD Life science Co.,LTD; Nanoject III, Drummond Scientific Company).

Three minutes following injection, to allow for adequate dispersal and absorption of the virus, the injector was removed from the animal; the incision site was closed and sutured. The mice received prophylactic analgesics before and after surgery (carprofen, 5 mg/kg, subcutaneous). Mice were at least eight weeks of age at the time of injection. With the exception of longitudinal observational studies on TetTox-treated animals, all animals were allowed at least 3 weeks to recover following surgery before initiating studies.

### Food intake studies

For acute food intake studies, animals on chow or 60%HFD (Research diets inc., D12492) were fasted four hours prior to onset of the dark cycle. Immediately prior to the onset of the dark cycle, animals were given an intraperitoneal injection of vehicle, clozapine-n-oxide (CNO, 1 mg/kg), liraglutide (400 ug/kg), or semaglutide (10 nmol/kg). A known amount of food was then returned, and remaining food was measured 2 and 4 hours later.

For chronic hM3Dq studies, mice were given saline for one to two days prior to injecting saline or CNO (IP, 1 mg/kg at 5 PM) followed by providing CNO (3.33 μg/ml) or vehicle in drinking water for the duration of treatment. Food intake and body weight were measured once daily at the onset of the dark cycle.

For chronic liraglutide studies, mice were administered vehicle (sodium phosphate 50 mM, sodium chloride 70 mM, 0.007% Tween-20, pH=8) for two days, followed by an increasing dosing paradigm over the first three days of 100 ug/kg, 300 ug/kg, and 500 ug/kg, after which the maximum dose was used throughout the treatment period. Injections were administered once daily, one hour prior to the onset of the dark cycle, and food intake and body weight were measured once daily at time of injection.

For chronic semaglutide studies, mice were administered the aforementioned vehicle for two days, followed by an increasing dosing paradigm over the first four days of 2.5 nmol/kg, 5 nmol/kg, 7.5 nmol/kg, and 10 nmol/kg, after which the maximum dose was used throughout the treatment period. Injections were administered once daily, one hour prior to the onset of the dark cycle, with food intake and body weight recorded at time of injection.

### Conditioned taste avoidance paradigm

For AP^Glp1r-Dq^ mice: Mice that were 8 weeks of age or older were individually housed in standard cages with free access to food and water. Mice were habituated to two water-containing bottles for 3-5 days until they learned to concentrate their daily water consumption. On the conditioning day, the mice received only saccharin (0.15%, 240931, Sigma) bottles. Following 30 min exposure to saccharin, mice were then injected intraperitoneally with the desired stimulus (vehicle control (0.9% NaCl), LiCl (126 mg/kg, 203637, Sigma) or CNO (1 mg/kg, C879476, Macklin). Access to the saccharin continued for an additional 1 h, followed by the return of normal water bottles. Two days later, each mouse received access to two water bottles (one containing 0.15% saccharine, the other containing water), and the amount of fluid ingested from each water bottle was measured.

For NTS^Glp1r-Dq^ mice: following an overnight fast, mice that were 8 weeks of age or older were conditioned with HFD (D012492, Research Diets) and paired with Saline, LiCl (126 mg/kg, 203637, Sigma) or CNO (1 mg/kg, 4936, Tocris Bioscience). The treatments were administered 30 min after exposure an extra one-hour access for HFD. On the post-conditioning day, fasted mice received access to both HFD and chow and the consumption of each was measured.

Liraglutide: Mice were singly housed with *ad libitum* access to two bottles containing water. For four days, mice received mock injections of vehicle, and initial bottle preference was identified. Mice were then water deprived overnight at the onset of the dark cycle of day 4, and on day 5 were given 60 minutes of free access to two bottles containing 0.15% saccharin in water. At 60 minutes, mice were administered liraglutide (400ug/kg), with continued access to saccharin for an additional 60 minutes (120 min. total). Bottles were then removed, and water was returned to mice at the onset of the dark cycle of the conditioning day. On the onset of the dark cycle of day 6, mice were water deprived overnight. On day 7, mice were given 2 hours access to one bottle filled with water, while the other bottle was filled with 0.15% saccharin. Preference was calculated as volume of saccharin consumed divided by total fluid consumed from both bottles.

### Gastric emptying

Mice were fasted for four hours prior to oral gavage of 20 mg/ml paracetamol in saline. Mice received an injection of liraglutide 400 ug/kg 1 hour prior to gavage. 15 minutes after gavage, we took one blood sample via tail vein collection. Blood was then spun at 10,000 RCF at 4° C for 10 minutes. Plasma was collected and frozen at -80°C. We used an Acetaminophen L3K detection kit (Sekure Chemistry) to detect plasma paracetamol concentrations via a Molecular Devices Spectramax m5, and reported concentration of plasma paracetamol as mg/dL.

### Tissue prep, cDNA amplification and library construction for 10x snRNA-seq

*Glp1r^Cre^* mice on the *Gt(ROSA)26Sor^tm5.1(CAG-Sun1-sfGFP)Nat^* background (Jax Catalog #021039) were euthanized using isoflurane and decapitated, the brain was subsequently removed from the skull and sectioned into 1 mm thick coronal slices using a brain matrix. The hypothalamus was dissected and flash frozen in liquid N_2_.

On the day of the experiment, frozen tissue from mixed male and female mice was homogenized in Lysis Buffer (EZ Prep Nuclei Kit, Sigma) with Protector RNAase Inhibitor (Sigma) and filtered through a 30mm MACS strainer (Myltenti). Strained samples were centrifuged at 500 rcf x 5 minutes and pelleted nuclei were resuspended in washed with wash buffer (10 mM Tris Buffer pH 8.0, 5 mM KCl, 12.5 mM MgCl_2_, 1% BSA with RNAse inhibitor). Nuclei were strained again and recentrifuged at 500 rcf x 5 minutes. Washed nuclei were resuspended in wash buffer with propidium iodide (PI, Sigma) and stained nuclei underwent FACS sorting on a MoFlo Astrios Cell Sorter. Double-labelled GFP+/PI+ nuclei were collected. Sorted nuclei were centrifuged at 100 rcf x 6 minutes and resuspended in wash buffer to obtain a concentration of 750 – 1200 nuclei/uL. RT mix was added to target ∼10,000 nuclei recovered and loaded onto the 10x Chromium Controller chip. The Chromium Single Cell 3’ Library and Gel Bead Kit v3, Chromium Chip B Single Cell kit and Chromium i7 Multiplex Kit were used for subsequent RT, cDNA amplification and library preparation as instructed by the manufacturer. Libraries were sequenced on an Illumina NovaSeq 6000 (pair-ended with read lengths of 150 nt).

### snRNA-seq data analysis

The FASTQ files were mapped to the appropriate genome (Ensembl GRCm38 or Rnor_6.0) using cellranger to generate count matrix files and data was analyzed in R. Genes expressed in at least 5 cells in were retained. Cells with at least 600 detected genes were retained. Doublets were scored using Scrublet^32^ clusters with a median doublet score > 0.3 or individual cells with scores >0.3 were removed.

The data was then normalized using scran^33^ and centered and scaled for each dataset independently and genes that were called variable by Seurat *FindVariableFeatures* were input to PCA. The top PCs were retained at the “elbow” of the scree plot (normally 15-30, depending on the dataset) and then used for dimension reduction using UMAP and clustering using the Seurat *FindNeighbors* and *FindClusters* functions. *FindClusters* was optimized for maximizing cluster consistency by varying the resolution parameter from 0.2 upward in steps of 0.2 until a maximal mean silhouette score was found. Clusters were then hierarchically ordered based on their Euclidean distance in PC space and ordered based on their position in the tree.

Cell types were identified by projecting labels from published single cell RNA-seq datasets^13–15^ using the Seurat CCA method. Neuron cluster names were chosen based on genes found in unbiased marker gene search (Seurat *FindMarkers*). Clusters without unique marker genes were labeled by their neurochemical identity (GABA or GLU) plus a number.

### Perfusion and Immunohistochemistry

At the conclusion of all experiments mice were deeply anesthetized with isoflurane and transcardially perfused with phosphate-buffered saline (PBS) followed by 10% buffered formalin. Brains were removed, placed in 10% buffered formalin for 4-24 hours, and dehydrated in 30% sucrose for at least 48 hours. Using a freezing microtome (Leica, Buffalo Grove, IL), brains were cut into 30-um sections and collected in four series.

For all immunohistochemistry procedures, sections were blocked for 60 minutes (PBS with 0.03% triton x-100, 3% normal donkey serum). The sections were incubated overnight (12-24 hours) at room temperature using primary antibodies (Rabbit anti-GLP-1R, #EPR21819 Abcam. 1:500; Rabbit anti-FOS, #2250, Cell Signaling Technology, 1:1000; mouse anti-CGRP, ab81887, 1:1000, Abcam; Chicken anti-GFP, GFP1020, Aves Laboratories, 1:1000;). Tissue-bound antibodies were detected with species-specific Alexa Fluor-488, 568 or 647 conjugated secondary antibodies (Invitrogen, Thermo Fisher, 1:250). Images were collected on an Olympus (Center Valley, PA) BX53F microscope.

For FOS counts, we counted detectable FOS positive soma in the AP of saline or liraglutide (400 ug/kg, IP) treated AP^Ctrl^ and AP^GLP1R-KO^) animals. Numbers of FOS positive soma per section in the AP were averaged and compared across groups.

## Acknowledgements

We thank members of the Qiu, Myers and Pers labs, along with Thomas Lutz and Christelle Le Foll and their lab members, and the many colleagues at Novo Nordisk who contributed helpful discussions to these studies. Thanks to Kelli Rule and Katrina Fox for breeding colony management. Thanks to Neil Bloc and Jae-Hoon Shin for technical assistance in initial gastric emptying experiments. Supported by NIH R01DK141495 (to MGM), a grant from Novo Nordisk, Inc (to TP and MGM), NNF18OC0033444 to the Center for Adipocyte Signaling to FS, and NSFC (NO. 32371212) to WQ.

## Author Contributions

WTY, WQ, and MGM conceived the project, designed experiments, and analyzed data. WTY, YW, GZ, SH, AR, SK, FS, MD, AJT, IW, TP, and WQ performed experiments and analyzed data. WQ and GZ performed all tetanus toxin, hM3Dq DREADD, and *Glp1r^Flox^*studies. AS, KR, RJS aided in project and experimental design and provided reagents. WTY and MGM wrote the manuscript with input from all authors. MGM and WQ acquired funding.

## Data availability

All data generated will be included in the manuscript and supplemental files.

---

**Supplemental Figure 1:**
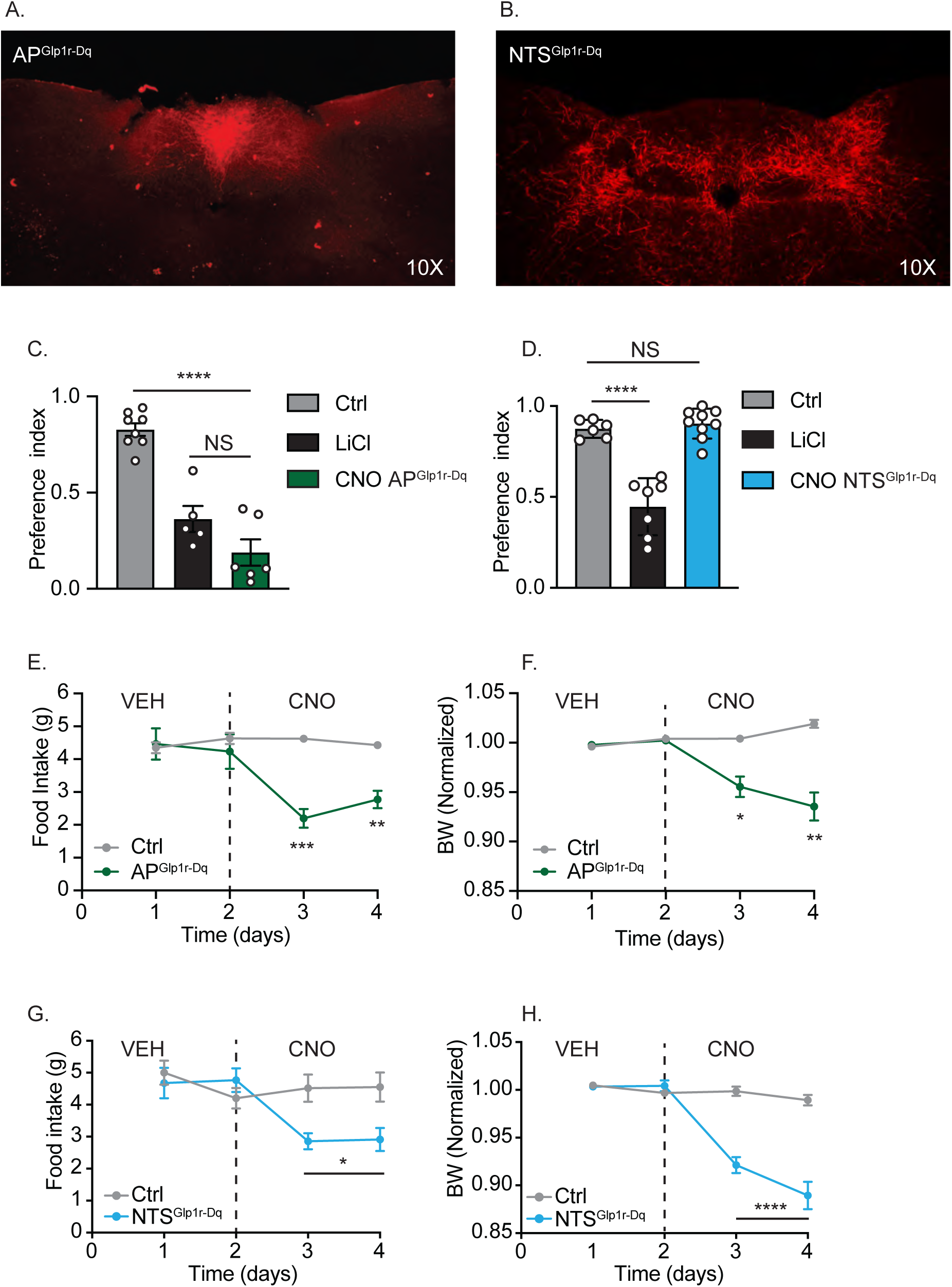
Activating AP^Glp1r^ or NTS^Glp1r^ neurons suppresses food intake, but only AP^Glp1r^ cells promote CTA formation. (A-B) Example images of mCherry-IR (red) in the DVC of AP^Glp1r-Dq^ (A) and NTS^Glp1r-Dq^ (B) mice (images acquired at 10x magnification). (C) Preference index for saccharin in AP^Glp1r-Dq^ mice treated with saline (Ctrl, n=8), LiCl (126 mg/kg) (n=5), or CNO (1mg/kg, n=6). (D) Preference index for HFD in NTS^Glp1r-Dq^ mice treated saline (CTRL, n=6), LiCl (126 mg/kg) (n=7), or CNO (1mg/kg, n=9). (E-F) Control (n=6) and AP^Glp1r-Dq^ (n=6) mice were given a loading dose of CNO (1mg/kg, IP) followed by 48-hour *ad libitum* access to CNO in a water bottle while monitoring food intake (E) and body weight (F) daily. (G-H) Control (n=6-8) and NTS^Glp1r-Dq^ (n=9-13) mice were given a loading dose of CNO (1mg/kg, IP) followed by 48-hour *ad libitum* access to CNO in a water bottle while monitoring food intake (G) and body weight (H) daily. Body weight in F, H are normalized to baseline food intake. All graphs: mean +/- SEM is shown. One way ANOVA with tukey’s test for C-D. Repeated measures two-way ANOVA with Sidaks multiple comparisons for E-H.. NS p>0.05, *p<0.05, **p<0.01, ***p<0.001, ****p<0.0001.

**Supplemental Figure 2:**
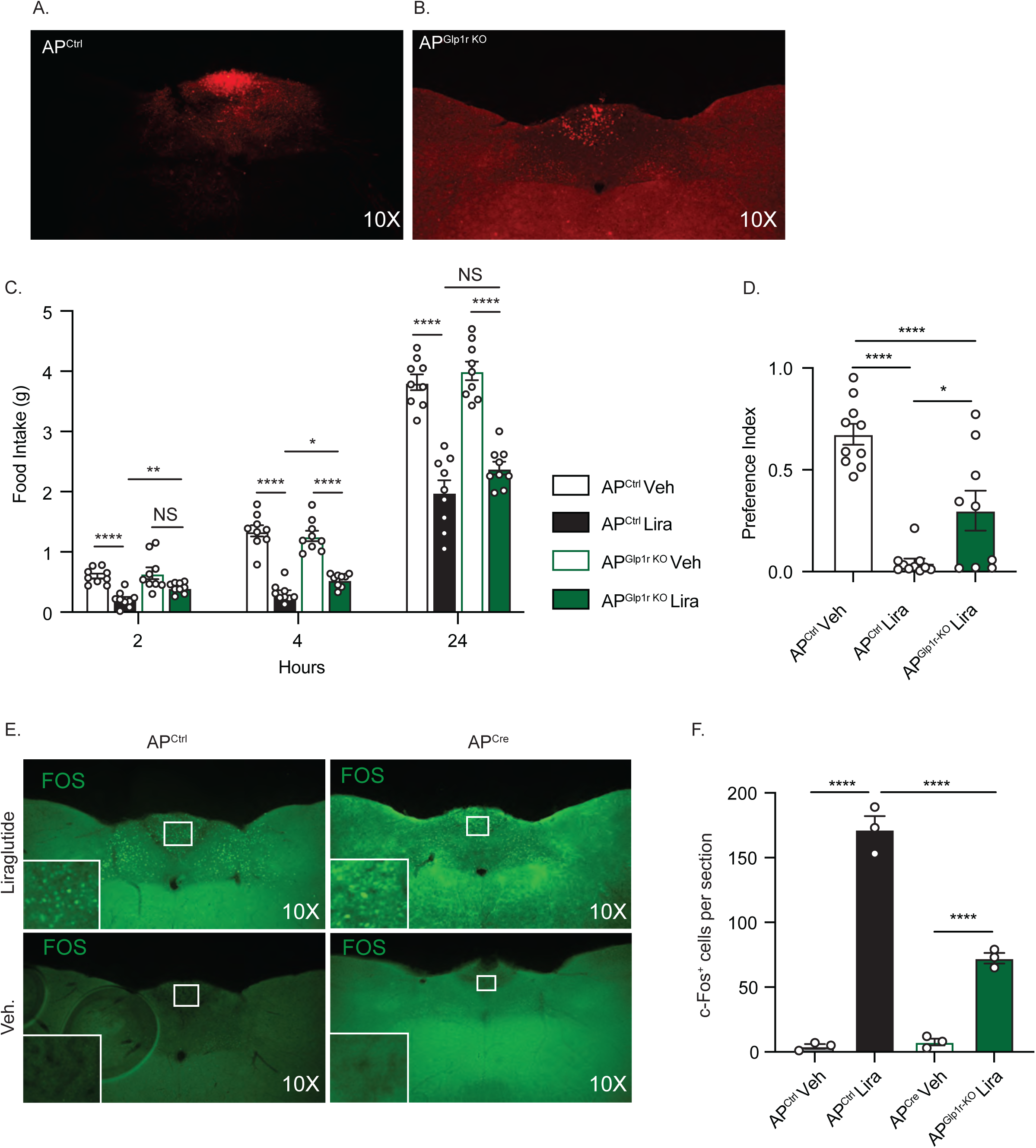
AP^Glp1r^ neurons contribute to the food intake and aversive responses to Lira. (A-B) Representative images of mCherry-IR (red) for AP^Ctrl^ and AP^Glp1r-KO^ mice (images acquired at 10X magnification). (C) Food intake beginning at the onset of the dark cycle for AP^Ctrl^ (n=9) and AP^Glp1r-KO^ (n=9) animals following vehicle (Veh) or Lira (400ug/kg, IP) injection. (D) Preference index calculated for saccharin in AP^Glp1r^ ^KO^ (n=9) or AP^Ctrl^ (n=10) animals treated with vehicle (Veh) or Lira (400 ug/kg, IP), as indicated. (E) Representative 10X images of FOS-IR (green) in the DVC of AP^Ctrl^ and AP^Glp1r-KO^ 90 minutes after vehicle and Lira (400 ug/kg) treatment. (F) Quantification of AP FOS-IR cells for animals treated as in (E) (n=3 per group). All graphs: mean +/- SEM shown. Two-way ANOVA w/Tukey’s post-hoc test for C, F. One-way ANOVA w/Tukey’s multiple comparisons for (D), NS p>0.05, *p<0.05, **p<0.01, ***p<0.001, ****p<0.0001.

**Supplemental Figure 3:**
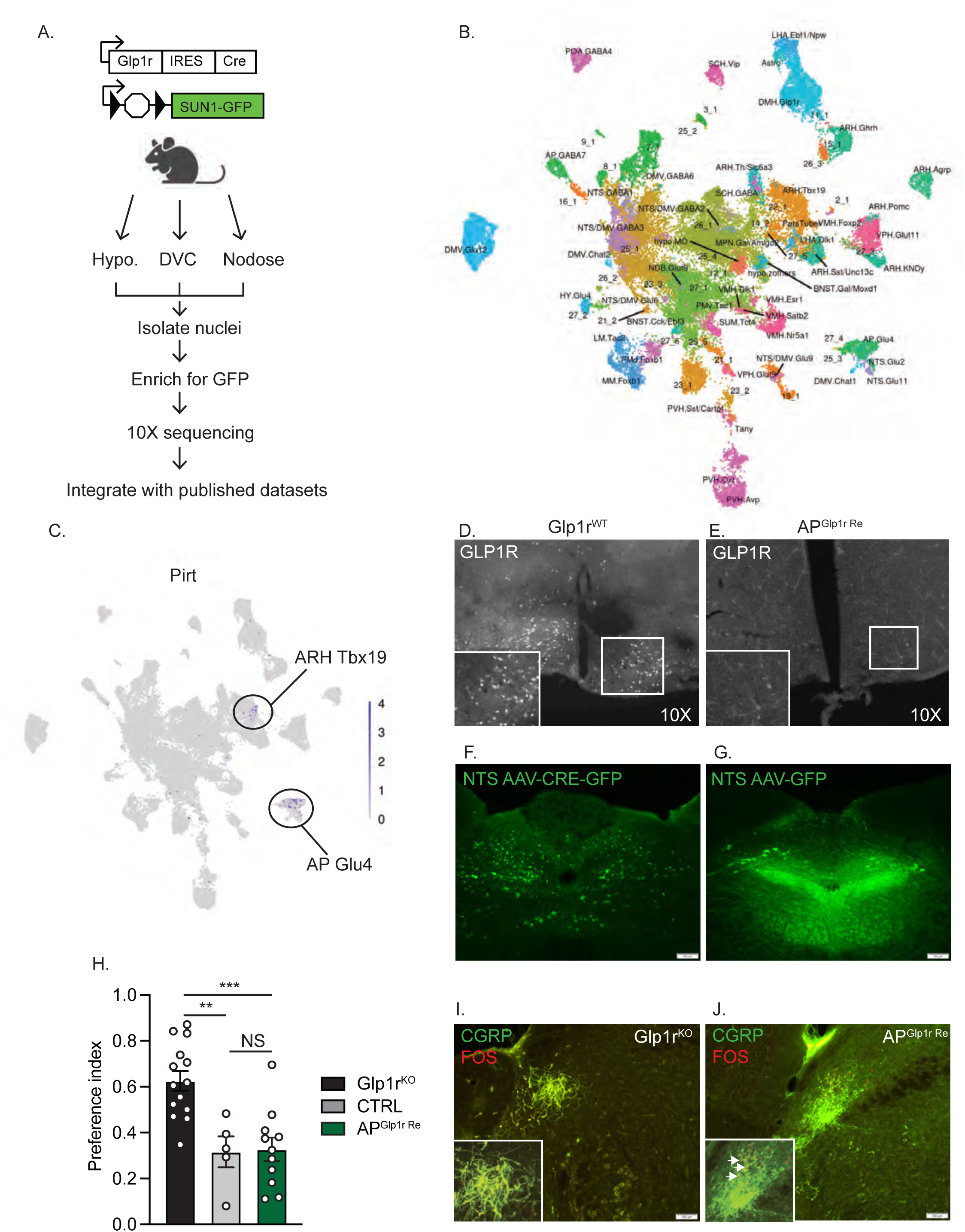
Identification of *Pirt* as marker gene for AP^Glp1r^ neurons. (A) Schematic diagram for snRNAseq analysis workflow. (B) UMAP clustering of snRNA-seq data from (A), with labeling of previously-identified and novel populations of *Glp1r* neurons. (C) UMAP showing *Pirt* expression (purple) across all clusters in (B), including labeling of *Pirt*-expressing populations. (D-E) Representative images showing GLP1R-IR (white) in the ARC of control (Glp1r^WT^) and *Pirt^Cre^;Glp1r^LoxTB/LoxTB^*(AP^Glp1r-Re^) animals (n=3 each, 10X magnification). (F-G) Example images of AAV-Cre-GFP or AAV-GFP injected into the NTS of *Glp1r^LSL/LSL^* mice Scale bar= 100um. (H) Preference index for saccharin in Glp1r^KO^ (n=14), Glp1r^WT^ (Ctrl; n=5), or AP^Glp1r-Re^ (n=11) mice treated with liraglutide (400ug/kg, IP). (I-J) Representative images showing CGRP-IR (green) and FOS-IR (red) in the l.PBN of Glp1r^KO^ (I) and AP^Glp1r-Re^ (J) animals treated with Sema (10 nmol/kg) for 2 hours. Scale bars=100 um. One-way ANOVA with tukeys (H). NS p>0.05, *p<0.05, **p<0.01, ***p<0.001, ****p<0.0001.

